# Adult thymus-derived cMaf^+^RORγt^+^ γδ T cells lack Scart2 chromatin accessibility and do not reach periphery

**DOI:** 10.1101/2023.02.20.529224

**Authors:** Tao Yang, Joana Barros-Martins, Anika Janssen, Ziqing Wang, Ximena León-Lara, Siegfried Weiss, Immo Prinz, Reinhold Förster, Sarina Ravens

## Abstract

T cell receptor (TCR) Vγ4^+^ expressing γ*δ* T cells can be divided into IFN-γ and IL-17-producing effector T cell subsets. A bias towards γ*δ*17 effector fate decisions is observed during early ontogeny. In contrast, the existence of Vγ4^+^ γ*δ*17 cells derived from adult thymus is still controversial. In the present work, we used a mouse model where T cells are exclusive generated within an adult thymus. Additionally, we employed single-cell chromatin state analysis from thymocytes of normal mice. A small, but considerable population of immature *Cd24*^+^ *Gzma*^+^ Vγ4 cells was found that exhibit molecular programs of γ*δ*17 cells. These adult thymus-derived immature *Cd24a*^+^ *cMaf*^+^ Vγ4 cells secrete small amounts of IL-17A and IL-17F. Interestingly, do not reach the periphery under steady-state conditions. Furthermore, *de novo* generated γ*δ*17-like cells from adult thymus lack transcriptional activity of the Scart2 encoding gene, suggesting that Scart2 is a distinct trait of fetal γ*δ* T cell precursors. Together, this study provides valuable insights into developmental traits of Vγ4 cells during adulthood and raises the question on signals suppressing the full maturation and/or thymic export of γ*δ*17-like cells within the adult thymus.

**Highlights:**

- Transcriptional and epigenetic profiling identifies developmental plasticity of *Gzma*^+^ *Cd24a*^+^ Vγ4 cells in adult thymus.
- Thymic c-Maf^+^ and RORγt^+^ Vγ4 T cells can be generated during adulthood, but do not reach the periphery under steady-state conditions.
- Innate CD44^high^CD45RB^neg^ γ*δ*17 cells are completely absent upon induction of T cell development during adulthood.
- Scart2 expression might be a key molecule to track developmental traits of fetal-derived γ*δ*17 cell precursors.

## Introduction

The functionality of γ*δ* T cells in mouse models of many diseases has been associated with the immediate production of two cytokines, interferon-γ (IFN-γ) and interleukin 17A (IL-17). Accordingly, they are divided into IL-17-producing (γ*δ*17) or IFNγ-producing (γ*δ*1) effector cells. These two groups further vary by variable gene segment usage of the T cell receptor (TCR) γ-chain and their ontogenetic origin.^1,2^ The majority of such cells, commit to γ*δ*1 or γ*δ*17 effector cell fate during fetal thymic development. However, only few of them leave the thymus as naïve cells. The γ*δ*1 type of cells include fetal-derived Vγ5^+^ dendritic epidermal T cells of the skin, Vγ7^+^ intraepithelial lymphocytes, and postnatal-derived Vγ1 and Vγ4 cells of lymphoid tissues. Expression of surface molecules CD27 and CD45RB is one common way to define γ*δ*1-committed cells.^3,4^

The large majority of γ*δ*17 cells expresses Vγ4^+^ or Vγ6^+^ (semi)invariant TCRs.^5^ Development preferably takes place during the embryonic period. Interestingly, functional commitment appears to occur prior to TCR rearrangement. ^33,6^ Within this time window, they develop from thymic CD24^+^ progenitors into IL-17 producers that can be associated with a CD44^high^CD45RB^neg^ surface phenotype in the periphery.^3^ Development of IL-17-producing Vγ6 cells, that are also expressing the Scart1 surface protein, is exclusively restricted to the embryonic thymus, where they dominate the γ*δ*17 cell pool.^3,6^ During ontogeny, they are followed by development of Vγ4 cells. They exhibit a commitment bias towards IL-17-producing effector cells initially.

After thymic egress, γ*δ*17 cells adapt as long-living cells to defined body locations. Both, Vγ4 and Vγ6 cells, display almost identical tissue-specific gene expression programs and functionalities.^7–10^ These IL-17-producing Vγ4 and Vγ6 cells, differ only in few minor features. Vγ4^+^ γ*δ*17 cells can additionally be defined by *Scart2* and *Cd9* surface expression,^9^ a higher TCR repertoire diversity^11^ and higher migration capabilities across tissues.^12^ Moreover, functionality of Vγ4 and Vγ6 cells appears to be controlled by TIM-3 or PD-1, respectively.^10^

Recent advances in next-generation sequencing technologies elucidated gene regulatory networks and TCR signaling molecules underlying functional differentiation of CD24^+^ γ*δ* cells (γ*δ*24 cells) into γ*δ*1 and γ*δ*17 cells. ^13–17^ These results now provide valid analytical resources. The γ*δ*1 cell subsets express key transcription factors of cytotoxicity and IFNγ-production, such as *Tbx21* (encoding T-bet) and *Eomes*, as well as natural killer receptors (NKRs). In addition, development of γ*δ*17 cells early in life is regulated by Notch signaling. Essential functional differentiation is also driven by a network of transcriptional regulators including *Maf, Sox13, Heb, Blk*, and *Rorc*.^14,18,19^ Since Vγ4^+^ cells are able to commit to one or the other effector pathway and keep the potential to rearrange Vγ4^+^ TCR during adult thymic development, we suggest a developmental plasticity at or after the appearance of the first perinatal Vγ4 T cell wave. However, detailed knowledge on molecular programs of progenitor or functional Vγ4 T cell subsets during adult thymic development is still lacking. Importantly, it was claimed that γ*δ*17 cells do not develop from adult hematopoietic stem cells and require an embryonic thymus to accomplish functional development. However, other studies provide evidence that naïve postnatal thymus derived Vγ4 cells subsets differentiate into IL-17 producers under certain pathological conditions.^10,20,21^ In the present work, transcriptional and epigenetic analysis Vγ4 cells from adult thymus and lymph nodes identify *de novo* generated c-Maf^+^ RORγt^+^ γ*δ*24 cells in adult thymus with retained capacity to co-secrete IL-17A and IL-17F. Most importantly, these *de novo* generated cMaf^+^ RORγt^+^ cells do not reach periphery and acquire a CD44^high^CD45RB^pos^ surface phenotype characterizing peripheral γ*δ*17 cells and lack transcriptional activity of the Scart2 encoding gene. Together, these data imply that innate CD44^high^ γ*δ*17 cells exclusively develop during the early time window of fetal thymus and supports the idea of inducible adult-thymus derived γ*δ* 17-like cells.

## Results

### Adult thymus γ*δ*24 cells exhibit transcriptional programs of γ*δ*1 and γ*δ*17 cells

T cells expressing a Vγ4^+^ γ-chain TCR produce either the cytokines IL-17 (γ*δ*17) or IFN-γ (γ*δ*1) (**Supp. Fig 1A**).^5^ The development of CD24^+^ Vγ4 cells is biased towards a higher generation of γ*δ*17 effector cells during the neonatal period as compared to later time points (**Fig. 1A**). We noted a small percentage of developing CD24^+^ Vγ4 cells that produce the cytokine IL-17A in adult thymus, though (**Fig. 1A**). To understand the underlying mechanisms of their functional commitment during adult thymus development and in peripheral lymphoid organs, droplet-based scRNAseq (single cell RNA sequencing) analysis was applied. To this end, adult thymus and peripheral lymph node (LN) Vγ4 T cells were isolated by fluorescent activated cell sortin. Transcriptomes of 13714 thymic and 3232 LN Vγ4 T cells were obtained after removing low-quality cells expressing < 200 genes and/or more than 10% mitochondrial genes. A non-linear dimensional reduction using uniform manifold approximation and projections (UMAP) of integrated thymic and LN Vγ4^+^ T cell transcriptomes was conducted. Based on differentially expressed gene (DEGs) analysis, twelve Clusters were identified (**Fig. 1B, Supp. Fig 1B**). Among the DEGs defining the respective Cluster, transcriptional regulators, cytokine and chemokine receptors as well as cell cycling genes were identified (**Fig. 1D**). Cells of Clusters c1, c2 and c12 represent exclusively thymus-derived Vγ4 cells and the majority of cells in c5-6 and c11 originates from peripheral LNs (**Fig. 1B-C, Suppl. Fig. 1C**).

**Fig. 1:**
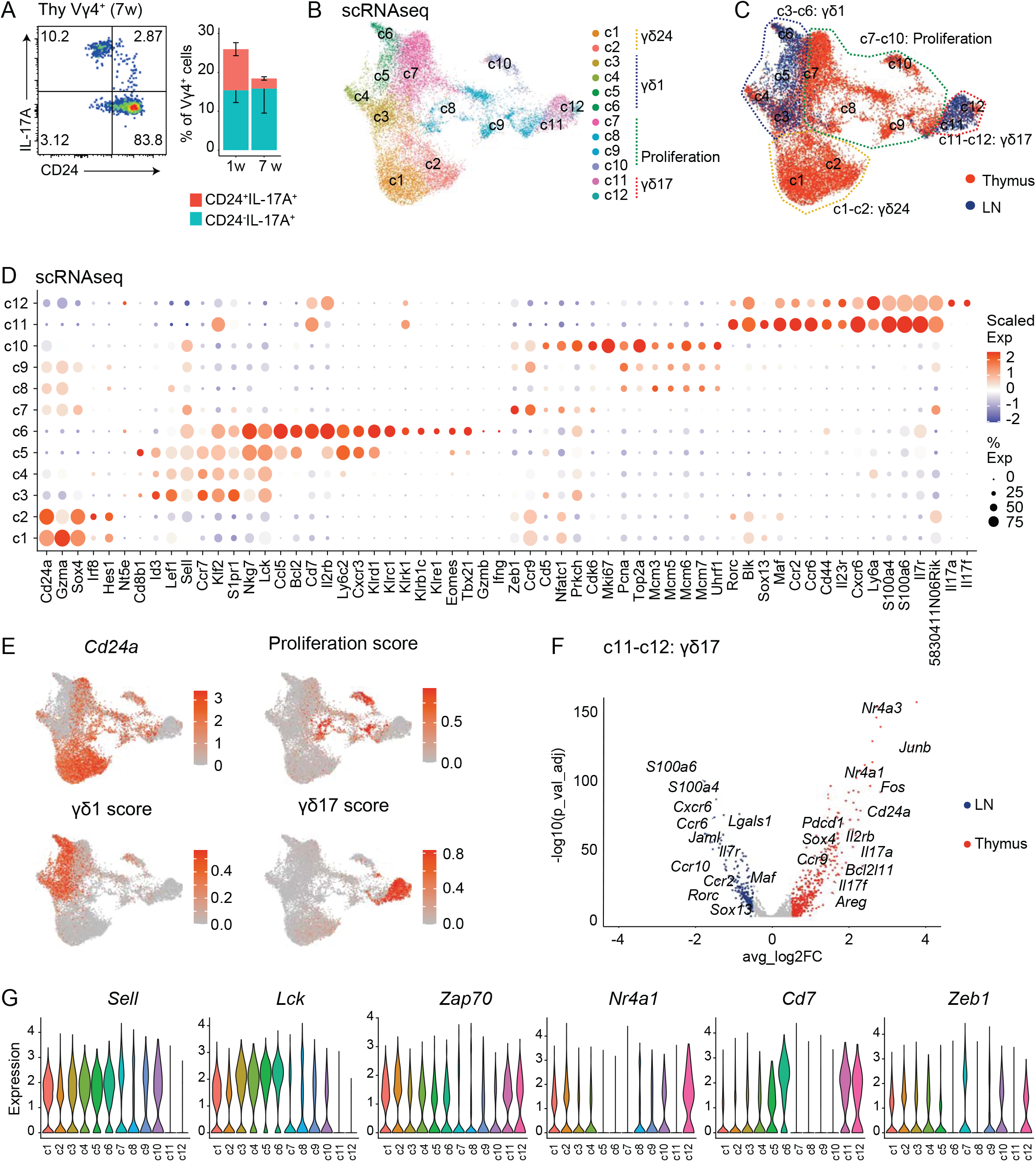
scRNAseq profiling of adult thymus and LN Vγ4 cells. (**A**) FACS analysis of IL-17A production relative to CD24 marker after ex vivo PMA and ionomycin stimulation in 1 week (N=10) and 3-7 weeks (N=6) old mice gated on thymus Vγ4^+^ cells. (**B-C**) Umap representation of adult thymus and LN Vγ4 cells scRNAseq data, colored by Cluster (**B**) and organ (**C**). Thymus and LN Vγ4 cells were sorted from 12 adult WT TcrdH2BeGFP mice. (**D**) Dot plots show the selected differentially expressed genes (DEGs) per Cluster. Gene expression values were scaled to a log2 fold change (logFC). Dots are colored by average logFC and sized by percentage of cells per Cluster that expressed this gene (% Exp). (**E**) Projection of the *Cd24a* expression level, as well as proliferation, γ*δ*1, and γ*δ*17 scores. The scores were computed with defined gene sets (proliferation: *Mki67, Cdk6, Pcna, Top2a, Mcm3, Mcm5, Mcm6;* γ*δ*1: *Tbx21, Eomes, Ifng, Gzmb, Lck, Cd5, Nfatc1, Prkch, Sell, Ccr7, Klf2, S1pr1, Nkg7*; *γδ17: Rorc, Maf, Blk, Sox13, Il17a, Il17f, Ccr2, Ccr6, Cxcr6, Cd44*); each cell is colored based on its individual score. (**F**) Volcano plot demonstrate DEGs of thymus and LN γ*δ*17 Vγ4 cells (c11 and c12). Upregulated DEGs are identified with log2FC > 0.5 and p_val_adj < 0.05. (**G**) Violin plots show normalized and log-transformed gene expression within each Cluster.

The phenotypic annotation of identified Clusters was based on key gene expression modules and DEGs that either relate to γ*δ*24, γ*δ*1, or γ*δ*17 cells (**Fig. 1E**).^9,13,17,22^ Accordingly, Clusters c1-c2 are defined as thymic *Cd24a*^*high*^ γ*δ*24 cells; thymic Clusters c7, c8-c10 as naïve cells with high proliferative potential (e.g., *Sell, Mik67, Cdk6*); pLN- and thymus-derived Clusters c3-c6 as naïve/effector γ*δ*1 cells (e.g., *Lck, Sell, Tbx1*); finally Clusters c11-c12 as γ*δ*17 cells (e.g., *Maf, Rorc, Il23r*) (**Fig. 1D-E**).

The γ*δ*1 cells are assigned to four Clusters, with c5-c6 cells exclusively originating from LNs (**Fig. 1C, Suppl. Fig. 1C**). The mixed Cluster c3-c6 are characterized by *Sell, Ccr7, S1pr1*, and *Lck* mRNA expression, which was previously described for naïve or γ*δ*1-committed cells.^13,22^ *Eomes* and *Tbx21* (encoding T-bet), key transcription factors regulating IFN-γ production, ^23–25^ are expressed in LN-derived Clusters c5-c6.

Specifically, genes encoding NKRs (e.g., *Klrc1, Klrk1* and *Klre1*) are expressed in *Eomes*^high^ and *Tbx21*^high^ LN-derived Cluster c6 (**Fig. 1D)**, which suggests their peripheral differentiation into cytotoxic effectors.^24^ The γ*δ*17 Clusters separate in LN-derived cells in c11 and exclusively thymus-derived cells in c12 (**Fig. 1B-C, Suppl. Fig. 1C**). To define differences between LN and thymus derived γ*δ*17 cells, a DEG comparison was conducted (**Fig. 1F**). Thymic γ*δ*17 Vγ4^+^ T cells display higher expression of *Bcl2* and *Bcl11* (anti-apoptosis), *Ccr9* (thymic migration) and the transcription factor *Sox4*, known to regulate *Sox13, Maf, Rorc* and *Blk* expression during development (**Fig. 1F**).^15^ Similarly, higher *Areg, Il17a* and *Il17f* gene expression of thymic Vγ4 cells was evident, equivalent to their Vγ6 counterparts.^9^ In contrast, enrichment of chemokine receptors (e.g., *Ccr2, Ccr6* and *Cxcr6*) in LN c11 cells suggests migratory capabilities.^12,26^ Moreover, LN cells express higher transcript levels of e.g., *S100a6, S100a8, Jaml* and *Lgals1*, which were previously assigned to fetal-derived γ*δ*17 cells in peripheral tissues.^7–9^

Next, γ*δ*24^+^ cells were characterized by high *Cd24a and Sox4* expression and can be divided into two Clusters. The γ*δ*24 cells of Cluster c1-2 may transition into highly proliferative *Cd24a*^*low*^ cells (c7-c10) that further express naïve T cell marker genes *Sell* and *Ccr9*, but not *Lck, Zap70* and *Nr4a1* (**Fig. 1D, Fig. 1G**). Within the proliferative *Cd24a*^+^ Cluster c7, high expression of the transcriptional regulator *Zeb1* is evident a zinc-finger E homeobox-binding transcription factor expressed in double-negative and double-positive thymocytes.^27^ The low expression of *Lck, Nr4a1* and *Cd7* in c7 suggests a modulation of TCR signals by *Zeb1* (**Fig. 1D, Fig. 1G**). Furthermore, detectable transcripts of transcription factors related to γ*δ*17 cell development, such as *Maf, Hes1, Blk, Sox13*^14,19,28,29^ were evident in γ*δ*24 Cluster 2 (**Fig. 1D**).

This is in line with the IL-17 production capability of CD24^+^ Vγ4 cells in adult thymus (**Fig. 1A**), Together, the transcriptomic data sets define gene expression programs of Vγ4 cells in LN and thymus and highlighted the separation of Vγ4 cells into γ*δ*24, γ*δ*1 and γ*δ*17 cells. The fact that adult thymus CD24^+^ Vγ4 cells are capable to produce IL-17 and express transcripts related to γ*δ*17 cell fate suggest a developmental plasticity at this stage.

### Granzyme A^+^ (Gzma) γ*δ*24 cells in adult thymus express c-Maf, but not RORγt

The transcription factor (TF) c-Maf is a key regulator of γ*δ*17 cell fate and known to induce *Rorc* expression, and thus IL-17 production capabilities.^14^ Accordingly, mature CD24^neg^ c-Maf^+^RORγt^+^ Vγ4 cells are present in the LNs (**Fig. 2A, left panel**). Importantly, flow cytometric analysis identified c-Maf^+^ and RORγt^+^ expression within adult thymus CD24^neg^ and CD24^+^ Vγ4 cells (**Fig. 2A**) similar to the IL-17 production capability of such cells (**Fig. 1A**). These observations are supported by detection of *Maf* transcripts within adult thymus γ*δ*24 Cluster c1-c2. They further displayed high levels of *Gzma* (encoding granzyme A) transcripts (**Fig. 2B**). *Gzma* expression was previously described for all developing γ*δ* T cells in fetal and adult thymus.^13,16^ Here we questioned about potential developmental trajectories of *Gzma*^+^ γ*δ*24 cells into γ*δ*1 and γ*δ*17 cells. Initial flow cytometric analysis of thymocytes confirmed that CD24^+^ Vγ4 cells might be able to produce GzmA.^16^ However, upon aging an increased abundance of CD24^neg^ T cells is observed (**Fig. 2C**).

**Fig. 2:**
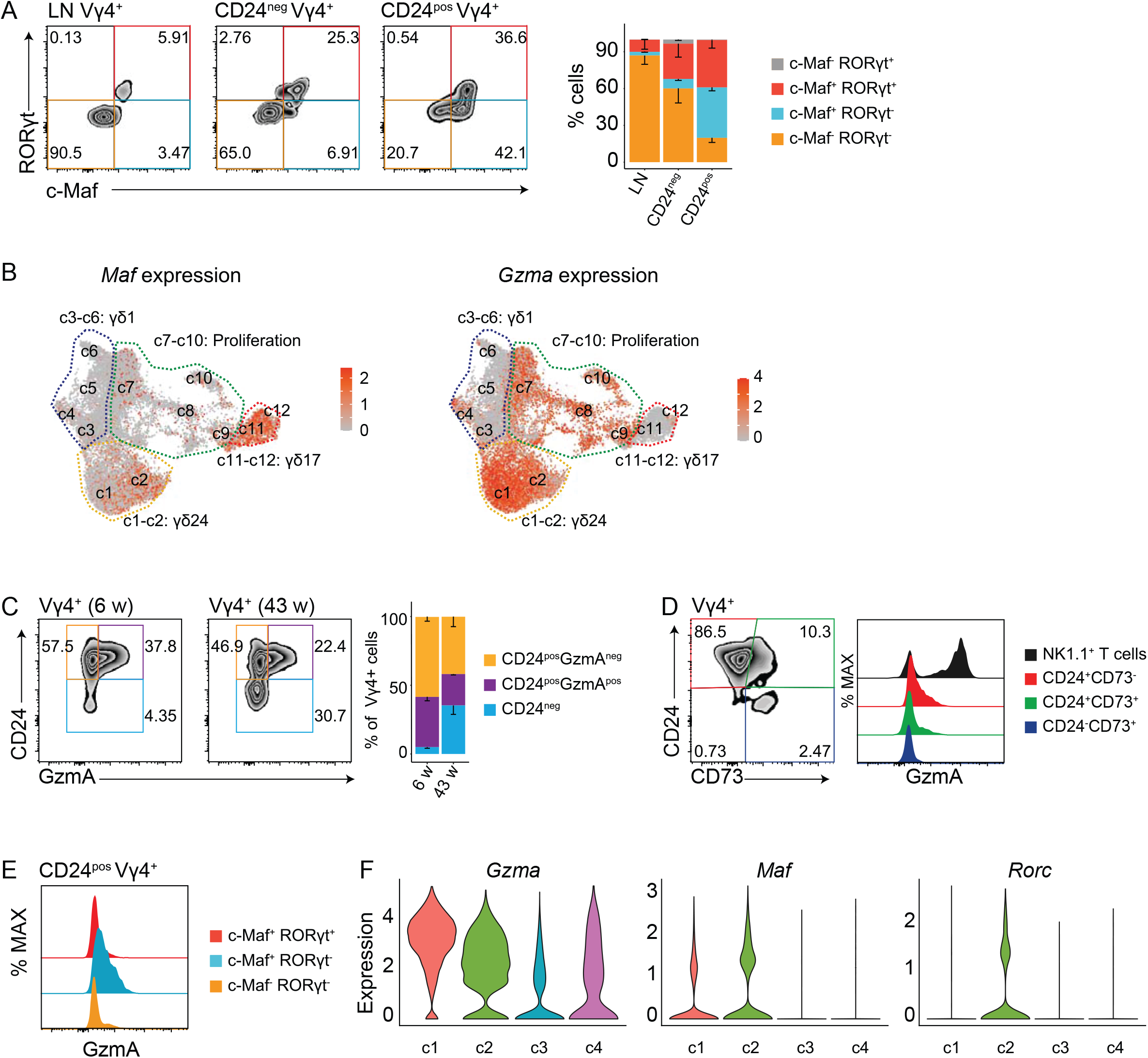
Identification of functionally uncommitted GzmA^+^ CD24^+^ Vγ4 T cells within thymus. (**A**) FACS analysis of the GZMA expression relative to RORγt and c-Maf transcription factor expression of LN, thymus CD24^-^, and CD24^+^ Vγ4 cells(N=3). (**B**) Umap displays *Maf* and *Gzma* expression. Each cell is colored based on its individual score. (**C-D**) FACS analysis of the GZMA expression relative to CD24 maturation marker in 6 weeks and 43 weeks old mice (N=3) (**C**) and CD24 and CD73 surface marker expression of thymus Vγ4^+^ cells (N=3) (**D**). (**E**) FACS plot showing the GzmA expression relative to RORγt and c-Maf expression gated on thymus CD24^+^Vγ4^+^ cells. (**F**) Violin plots show normalized and log-transformed gene expression within each Cluster, colored by Cluster (c1-c4).

The acquisition of effector cell fate of γ*δ* T cells during development can be monitored by surface expression of CD24 and CD73, i.e. the transition of CD24^+^ cells from the CD24^+^CD73^+^ double positive stage towards CD24^neg^CD73^+^ cells (**Fig. 2D**).^30^ Of note, GzmA production was highest in CD24^+^CD73^neg^ Vγ4 cells, but never reached the level of Nk1.1^+^ αβ NKT cells (**Fig. 2D, Supp Fig 2A**). Moreover, among all CD24^+^ Vγ4 T cells c-Maf^+^ RORγt^neg^ cells expressed the highest amount of GzmA (**Fig. 2E**). Together with varying gene expression levels of *Rorc, cMaf* and *Gzma* in γ*δ*24 Clusters c1-4 (**Fig. 2F**), these results point to a loss of GzmA expression upon functional commitment into c-Maf^+^RORγt^+^ cells (Cluster c2). In line, GzmA expression is completely undetectable in all LN γ*δ* cells (**Supp. Fig 2B**). Altogether, CD24^+^ thymic Vγ4 cells are capable to produce small amounts of GzmA that declines with preceding development into functional subsets. Moreover, combined scNGS and flow cytometric analyses reveal that adult thymus CD24^+^ Vγ4 cells express key transcription factors characteristic for IL-17 producers.

### Epigenetic profiling indicates developmental plasticity of CD24^+^ Vγ4 cells in adult thymus

Next, we wondered about the developmental potential of adult thymus γ*δ*24 cells. Single-cell ATACseq monitors the chromatin accessibility of promoter and gene coding regions within individual cells. As chromatin accessibility is taking place prior to gene expression, it can be applied as a predictive tool of transcriptional activity and developmental traits. Employing the same strategy as for scRNAseq, the chromatin state of thymic and LN Vγ4 T cells was determined by scATACseq. After removing low-quality cells with transcriptional start site (TSS) enrichment score < 2.5 and sequencing depth score <60, a total of 7031 thymic and 4736 LN cells were subjected to integrated analysis. Re-unsupervised clustering with adjustment by differential accessible gene region (DAR) analysis among neighboring Clusters identified 10 Clusters (c1-10) with a partial overlap of thymic and LN cells (**Fig. 3A-C, Suppl. Fig 3A**). The majority of DARs locates in promoter regions within 3kb of the nearest transcriptional start site confirming that the data set robustly detects accessible gene regions (**Suppl. Fig 3B**). Coverage plots of selected genes (e.g., *Il23r, Eomes* and *Ifng*) indicates data robustness as there is an accumulation of ATAC peaks within the promoter and gene body region (**Suppl. Fig 3C**).

**Fig. 3:**
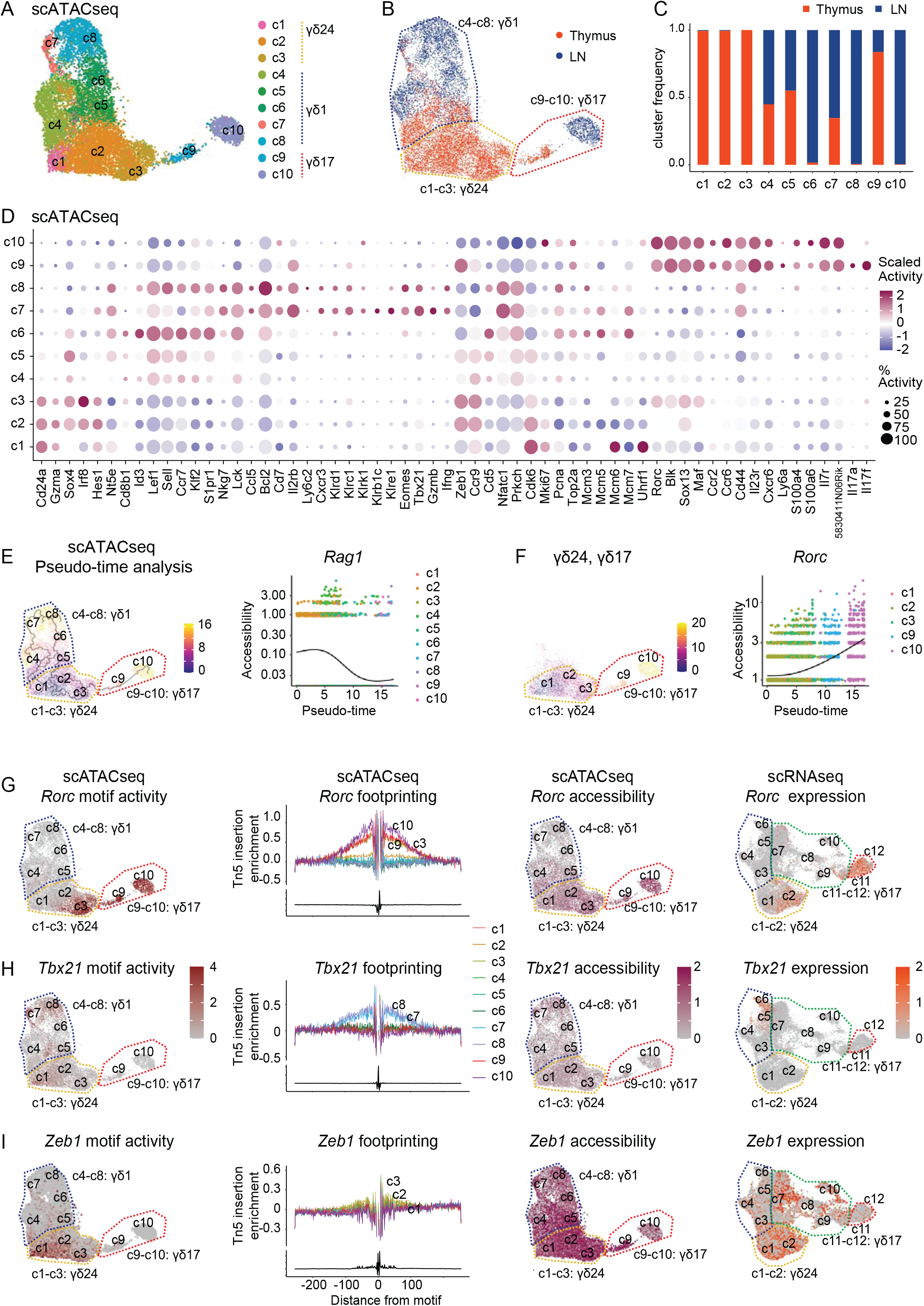
scATACseq profiling of adult thymus and LN Vγ4 cells. (**A-B**) Umap representation of adult thymus and LN Vγ4 cells scATACseq, colored by Cluster (**A**) and organ (**B**). Thymus and LN Vγ4 cells were sorted from 25 adult TcrdH2BeGFP mice. (**C**) Bar plot reveals fractions of absolute cell numbers from thymus and LN Vγ4 scATACseq data that contributed to c1 to c10. (**D**) Dot plots show selected differential accessible genes (DAGs) (gene body ± 2 kb) of each Cluster. Gene accessibility values were scaled to a log2 fold change (logFC). Dots are colored by average logFC and sized by percentage of cells per Cluster that open the accessibility (% Activity). (**E**) Left: Umap shows the pseudo-time analysis of Vγ4 cell scATACseq. Each cell is colored based on its individual pseudo-time score. Right: *Rag1* gene accessibility dynamics along the pseudo-time trajectory. (**F**)Left: Umap displaying the pseudo-time trajectory from the trait: γ*δ*24 (c1-c3) to γ*δ*17 (c9-c10). Right: *Rorc* gene accessibility dynamics along the pseudo-time trajectory. (**G-I**) Umaps highlighting *Rorc* (**G**), *Tbx21* (**H**) and *Zeb1* (**I**) motif activity (the 1st column), gene accessibility (the 3rd column), and gene expression level (scRNAseq, the 4th column). Each cell is colored based on its individual score. TF footprints (average scATACseq around predicted binding sites) of respect TFs for each Cluster (the 2nd column), colored by Cluster.

Based on differential accessibility of genes relating to T cell differentiation and functionality, c1-c10 can be divided into three main groups (**Fig. 3D, Suppl. Fig 3D**). Clusters c4-8 were annotated with γ*δ*1 T cells since they exerted high gene accessibility of *Cd27, Lef1* and genes related to TCR-dependent functional commitment, such as *Lck, Prkch, Cd5* and *Nfatc1* (**Fig. 3D)**. Similar to transcriptomic profiles, two largely LN-derived γ*δ*1 Clusters (c7-c8) displayed high gene accessibility of *Eomes, Tbx21*, NKR genes and the effector molecules *Gzmb* and *Ifng*.

For γ*δ*17 T cells, the thymus-biased Cluster c9 and LN-restricted Cluster c10 are separated from all other Clusters. They are characterized by high gene accessibility of γ*δ*17 lineage markers (e.g., *Rorc, Maf, Ccr2, Ccr6, Cd44, Il23r)*. A few details light up (**Fig. 3D**): in line with transcriptional profiles in Figure 1, the *Il17a* and *Il17f* gene regions were open in thymus, but not in cells from LN. Similar to Vγ6^+^ γ*δ*17 cells,^9^ such LN Vγ4^+^ γ*δ*17 cells have a higher proliferative potential than the thymus derived cells.

Clusters c1-c3, annotated as γ*δ*24 cells, are exclusively thymus-derived and display high gene accessibility of *Cd24a, Sox4, Hes1, Ccr9, Zeb1* and *Gzma*, but also of cycling genes (e.g., *Cdk6, Mki67* and *Mcm6*) (**Fig. 3D**). Importantly, chromatin regions of genes related to TCR-driven activation/differentiation, e.g., *Lck* or *Prkch*, are not accessible in γ*δ*24 cells (**Suppl. Fig 3E**). Open chromatin of *cMaf, Rorc* and *Blk* gene loci within γ*δ*24 cells in c3, but no the other γ*δ*24 Clusters, as well as gene accessibility of *Zeb1* within thymus γ*δ*17 cells (c9) proposes that a fraction of these cells may have the potential to become IL-17-producing cells or recently completed development in adult thymus. Next, we aimed at predicting potential developmental trajectories of *Gzma*^+^ γ*δ*24 cells into effector cell subsets. Therefore, a pseudo-time analysis was conducted on the scATACseq profiles of thymic and LN Vγ4 cells (**Fig. 3E**). The thymic Cluster c2 with highest *Cd24a* gene accessibility was set as the root node of pseudo-time. The *Cd24a, Gzma* and *Rag1* chromatin accessibility are decreasing along pseudo-time, confirming robustness of the method in predicting T cell differentiation pathways (**Fig. 3E, Suppl. Fig. 4A**). Specifically, the UMAP in **Fig. 3E** suggests two developmental traits of *Cd24a*^high^ Vγ4 cells. One trait is the flow of cell state transitions from thymic *Cd24a*^high^ Clusters (c1-c3), to thymic γ*δ*17 Cluster (c9), and LN γ*δ*17 Cluster (c10) (**Fig. 3F**). The accessibility of TF genes *Rorc and Maf* for effector γ*δ*17 cells, as well as markers *Ccr2* and *Ccr6* for γ*δ*17is increasing across pseudo-time from thymuic γ*δ*24 Clusters c1-c3 to γ*δ*17 Cluster c9 and LN γ*δ*17 c10 (**Fig. 3F, Suppl. Fig. 4B**). Notably, thymus γ*δ*17 cells of c9 are closer related to γ*δ*24 c3 cells than to LN-derived γ*δ*17 cells in c10 (**Fig. 3F, Suppl. Fig. 4B**). The LN c10 Cluster has the highest chromatin accessibility of γ*δ*17 marker genes across Clusters (e.g., *Rorc, Maf, Ccr2, Ccr6, Cxcr6, S100a4*, and *S100a6*) (**Fig. 3F, Suppl. 4B**).

**Figure 4:**
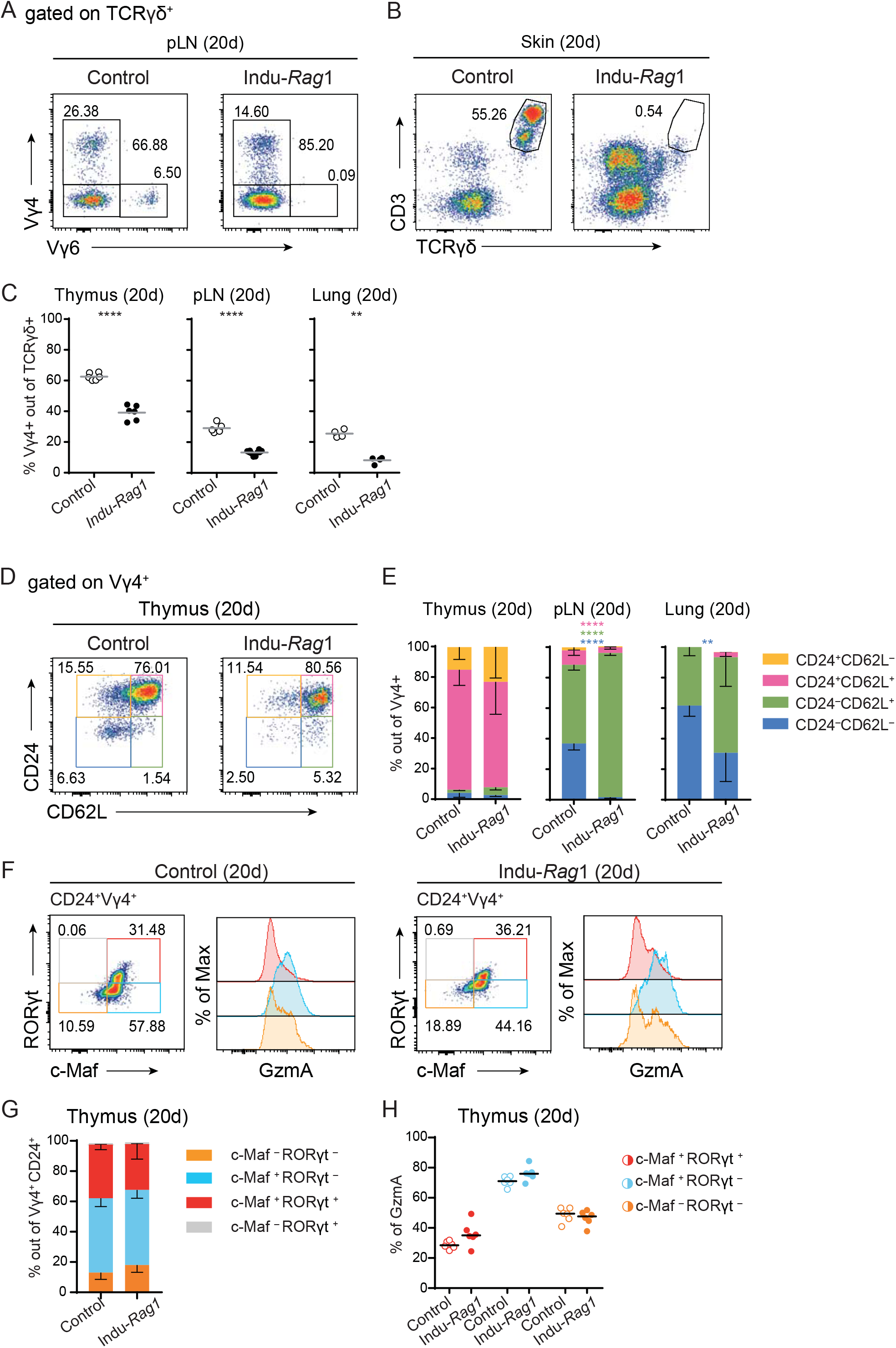
Exclusive development in adult thymus generates c-Maf^+^ RORγt^+^ Vγ4 T cells. (**A-B**) Analysis of Indu-*Rag1* peripheral lymph nodes (pLN) (**A**) and skin (**B**) for neonatal derived Vγ6^+^ and Vγ5^+^ γ*δ* T cells. (**C**) Quantification of flow cytometry analysis of Vγ4 usage in thymus, peripheral Lymph Nodes (pLN) and Lung of 20 days old controls and Indu-*Rag1* mice 20 days post induction. Gating strategy (**D**) and quantification (**E**) of flow cytometry analysis of CD24 and CD62L expression in Vγ4 γ*δ* T cells from thymus, pLNs and lung of 20 days old control mice and Indu-*Rag1* 20 days after induction of the *Rag1* gene. (**F**) Gating strategy of RORγt and c-Maf transcription factor expression in thymic CD24^+^Vγ4^+^ cells and GzmA expression relative to RORγt and c-Maf expressing populations in 20 days old controls and Indu-*Rag1* mice 20 days post induction. (**G**) Frequency of RORγt and c-Maf expressing populations out of CD24^+^Vγ4^+^ in 20 days old control mice and Indu-*Rag1* mice 20 days post induction. (**H**) Frequency of GzmA expression relative to RORγt and c-Maf expressing cells in 20 days old control mice (open circles) and Indu-*Rag1* mice 20 days post induction (filled circles). Data is representative of 4 different experiments with n=6 for 20 days old thymus from control and n=6 from Indu-*Rag1* mice, n=5 for 20 days old pLNs from control and n=10 from Indu-*Rag1* mice and n=4 for 20 days lung from control and n=4 from Indu-*Rag1* mice (C, E, G, H).

Another predicted trait starts from the thymus *Cd24a*^high^ Clusters (c1-c2), to the mixed γ*δ*1 Clusters (c4-c5), and ends at LN γ*δ*1 Clusters (c6-c8) (**Suppl. Fig. 4C**). In that case, the chromatin accessibility of γ*δ*1 marker genes is increasing (e.g., *Tbx21, Klrd1, Ifng, Lck, Eomes, Id3, Lef1, Cd7, Sell, Klf2, S1pr1, Nkg7, Ccl5, Il2rb* and *Klrc1*) (**Suppl. Fig. 4C**). Moreover, the accessibility of TCR signaling genes (e.g., *Lck, Zap70*, and *Cd7*) is increasing while the TCR signaling repressor *Zeb1* is decreasing (**Suppl. Fig. 4D**). TFs are key determinants of cell fate and their activity may relate to developmental traits. One advantage of scATACseq is to predict the TF activity of individual cells based on the presence of TF binding sites within open chromatin regions. It should support the pseudo-time analysis. The TF *Rorc* drives IL-17 production of T cells.^14,31^ The UMAP in **Fig. 3G** shows *Rorc* TF motif enrichment in γ*δ*17 Cluster c9 and c10, and thymic γ*δ*24 Cluster c3. Furthermore, examination of Tn5 integration events surrounding *Rorc* motifs in the TF footprints revealed a strong integration event flanking the TF motif in c3 as well as c9-c10. Results are confimed by increased *Rorc* chromatin accessibility and increased *Rorc* transcription (**Fig. 3G**). Moreover, binding patterns of T-bet (encoded by *Tbx21*) and *Eomes*, key TFs for IFNγ producing and cytotoxic γ*δ* T cells,^24^ were determined. In contrast to the detected RORγt motif activity in γ*δ*24 cells, *Tbx21* and *Eomes* motif activity, chromatin accessibility, as well as transcriptional activity was mainly detected in γ*δ*24^neg^ LN-derived γ*δ*1 Clusters (**Fig. 3H, Suppl. Fig. 4E)**. Of note, the motif activity and chromatin accessibility of transcriptional regulator *Zeb1* was enriched in γ*δ*24 cells and declined in γ*δ*1 Clusters (**Fig. 3I**).

In sum, the scATACseq provides a resource to determine gene regulatory programs of developing and mature Vγ4 cells in thymus and LN. The developmental trajectory analysis and presence of key TFs that govern γ*δ*17 differentiation proposes a considerable developmental plasticity within adult thymus γ*δ*24 cells. We speculate that γ*δ*24 cells in thymus, which harbor molecular programs of γ*δ*17 cells either (i) originated from the perinatal time window and did not fulfill complete differentiation yet or (ii) derive from *de novo* development of precursors in adult thymus. Their potential to mature into functional IL-17 producers is unclear.^6,20,21,32^

### c-Maf^+^ RORγt^+^ Vγ4 cells are generated within an adult thymus, but do not become CD44^high^CD45RB^neg^ γ*δ*17 cells

We thought to re-investigate the development of Vγ4 cells, exclusively generated from adult thymic precursor cells.^6^ For this purpose, we employed a previously established mouse model (named Indu-*Rag1*) where tamoxifen application rescues the inversion and inactivation of the *recombination-activating gene 1* (*Rag1*). This results in activation of T cell development within the thymus. Thus, γ*δ* T cells under these conditions arise from adult thymic precursor cells.^6,33^ At 20 days after *Rag1* gene activation, γ*δ* T cells were studied. Initial flow cytometric analysis verified the generation of Vγ1 and Vγ4 cells, but not Vγ6 cells, within Indu-Rag1 mice (**Fig. 4A, Suppl. Fig 5A-B**).^6,34^ The absence ofγ*δ* T cells in the Indu-*Rag1* skin was confirmed (**Fig 4B**) ^6^. Moreover, in line with a previous report,^32^ lung Vγ1 and Vγ4 cells believed to solely develop from adult thymus precursors were generated in such mice (**Suppl. Fig 5A-B**). Interestingly, 20 days post-induction Indu-*Rag*1 mice had a significant deficit of Vγ4 cells in both the thymus and peripheral organs as compared to control mice 20 days of age (**Fig 4C**).

During thymic development, γ*δ*24 T cells transition via the CD24^+^CD62L^+^ double positive stage into naïve CD24^neg^CD62L^pos^ cells and/or acquire a CD24^neg^CD62L^neg^ effector state (**Fig 4D**). Although thymic Indu-*Rag1* Vγ4 cells did not present any apparent differences to their WT counterparts (**Fig 4E - left panel**), Indu-*Rag1* Vγ4 cells in pLNs were predominantly composed of naive CD24^neg^CD62L^+^ cells that seem to undergo final maturation within tissues (**Fig 4E – centre and right panels**). On the other hand, Indu-*Rag1* Vγ1 cells exhibited no differences in lung compared to WT mice (**Suppl. Fig 5C – right panel**). However, pLN and to lower extent thymus Indu-*Rag1* Vγ1 cells mimicked the phenotype that was observed for Vγ4 cells 20 days post induction (**Suppl. Fig 5C – left and centre panels**).

Since our single-cell NGS identified adult γ*δ*24 cells that harbor molecular programs of γ*δ*17 cells (**Fig 1 and Fig 3**), we investigated expression of c-Maf and RORγt at 20 days post *Rag*1 activation. Theabundance of γ*δ*24 T cells expressing c-Maf^+^ and c-Maf^+^ RORγt^+^ was highly similar between WT and Indu-*Rag1* thymus samples (**Fig 4 F-G**). Similarly, no difference in GzmA production was evident among Indu-*Rag1* and WT γ*δ*24 cells at the respective cell states (**Fig 4H and Suppl. Fig 5D**).

In peripheral lymphoid organs and tissues, the surface molecules CD44 and CD45RB allow a segregation of γ*δ* cells into CD44^high^ CD45RB^neg^ γ*δ*17 and CD44^int^ CD45RB^+^ γ*δ*1 subsets, respectively.^3^ The presence of such CD44^high^ CD45RB^neg^ Vγ4 cells in pLNs (approx. 35%) and lung (approx. 60%) in WT control samples was confirmed.^3^ Importantly, CD44^high^CD45RB^neg^ Vγ4 cells were virtually absent in Indu-*Rag1* pLNs and lungs. This was supported by the high abundance of CD62L^+^ cells in Indu-*Rag1* pLN samples (**Fig 5A-B**).

**Figure 5:**
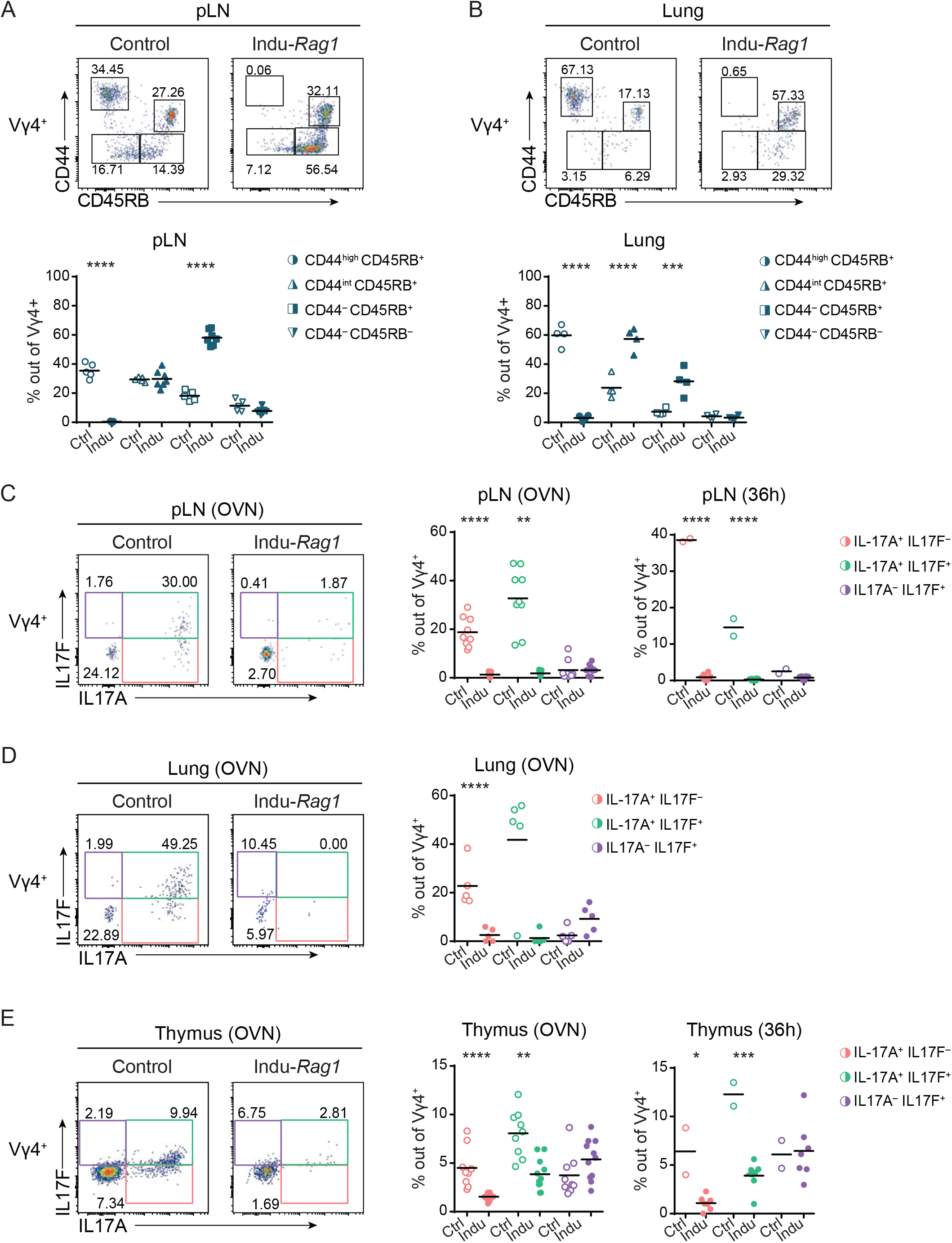
Adult thymus-derived cMaf^+^RORγt^+^ Vγ4 T cells have impaired functionality of IL-17A/F production and don’t reach the periphery. (**A-B**) Gating strategy (top) and quantification (bottom) of flow cytometry analysis of CD44 and CD45RB expression in peripheral Lymph Nodes (pLN) (**A**) and Lung (**B**) of 20 days old controls and Indu-*Rag1* mice 20 days. (**C-E**) Flow cytometry analysis of *ex vivo* enriched γ*δ* T cells suspensions from pLN (**C**), lung (**D**) and thymus (**E**). Cells from 20 days old controls and Indu-*Rag1* mice 20 days were stimulated with IL-1β and IL-23 for over-night or 36h and restimulated with PMA plus ionomycin for 4h with Brefeldin A before analysis. (**A-B**) Data is representative of 3 different experiments with n=5 for 20 days old thymus from control and n=7 from Indu-*Rag1* mice in (A), n=4 for 20 days old thymus from control and n=4 from Indu-*Rag1* mice in (B). (**C-E**) Data is representative of 3 different experiments with n=9 for 20 days old pLNs from control and n=11 from Indu-*Rag1* mice in (C), n=5 for 20 days old thymus from control and n=5 from Indu-*Rag1* mice in (D) and n=9 for 20 days old pLNs from control and n=11 from Indu-*Rag1* mice in (E).

Next, we investigated in the functionality of c-Maf^+^ RORγt^+^ Vγ4 cells generated exclusively in the adult thymus. Therefore, we isolated γ*δ* T cells from WT and Indu-*Rag1* thymi, pLNs and lungs and stimulated them *ex vivo* with the innate cytokines IL-1ß and IL-23 overnight or for 36h. In agreement with previous data,^6^ IL-17A production of Indu-*Rag1* Vγ4 cells within pLNs and lungs was significantly impaired as compared to controls (**Fig 5C-D**). Notably, a small, but visible proportion of isolated Vγ4 cells within Indu-*Rag1* thymic samples was able to co-secrete IL17A and IL-17F after overnight and 36h stimulation (**Fig 5E**).

Together, Vγ4^+^ γ*δ* T cells that express c-Maf and RORγt with the potential to co-secrete small amounts of IL-17A and IL-17F are newly generated within the adult thymus. However, these cells may follow different maturation traits compared to their CD44^high^ CD45RB^neg^ fetal-derived γ*δ*17 counterparts and do not reach the periphery at *steady-state*.

### Adult thymus-derived c-Maf^+^RORγt^+^ γ*δ*24 cells lack *Scart2* gene accessibility

To determine potential underlying molecular programs that may explain lower IL-17A and IL-17F production capability of adult-thymus derived c-Maf^+^RORγt^+^ γ*δ* T cells and their absence in the periphery, scATACseq of isolated Vγ4 cells from Indu-*Rag1* thymocytes was performed. Data were compared to epigenetic profiles of *TcrdH2BeGFP* reporter mice (WT) by integrated analysis (**Fig. 6A**). As represented in the UMAPs, in total seven Clusters were identified, relating to γ*δ*1 or γ*δ*17 cells (**Fig. 6A-C, Suppl. Fig 6A-C**). The bar plot confirms that cells from WT or Indu-*Rag1* thymus contribute to all identified Clusters (**Fig. 6C**). We only noted a higher abundance of γ*δ*24 cells in c2, a slight increase of γ*δ*1 cells (c6 and c7), and fewer γ*δ*17 cells (c3-4) in Indu-*Rag1* thymus (**Fig. 6C**). Genome feature plots illustrate a similar gene accessibility of *Cd24* and *Gzma* among WT and Indu-*Rag1* thymocytes in the respective Cluster (**Fig. 6D**). Differential accessible genes (DAGs) analysis among WT and Indu-*Rag1* thymus Vγ4 cells identified only 8 DAGs (log2FC > 0.5) or 282 DAGs (log2FC >0.25), respectively (**Fig. 6E, Supp. Fig 6D**). Indu-*Rag1* cells in Cluster c4 show a high γ*δ*17 cell gene accessibility score and high γ*δ*17-associated genes like *Sox13, Sox4, Blk* and *Maf*, as well as motif activity of the TF RORγt, which is similar to WT cells (**Fig. 6F, Suppl. Fig 6C**). The Cluster c7 display the highest γ*δ*1 score, as well as TF motif activity of *Tbx21* and *Eomes* (**Fig. 6F, Suppl Fig. 6C**). Together, epigenetic profiles among Indu-*Rag1* and WT thymus cells display only small nuanced differences.

**Fig. 6:**
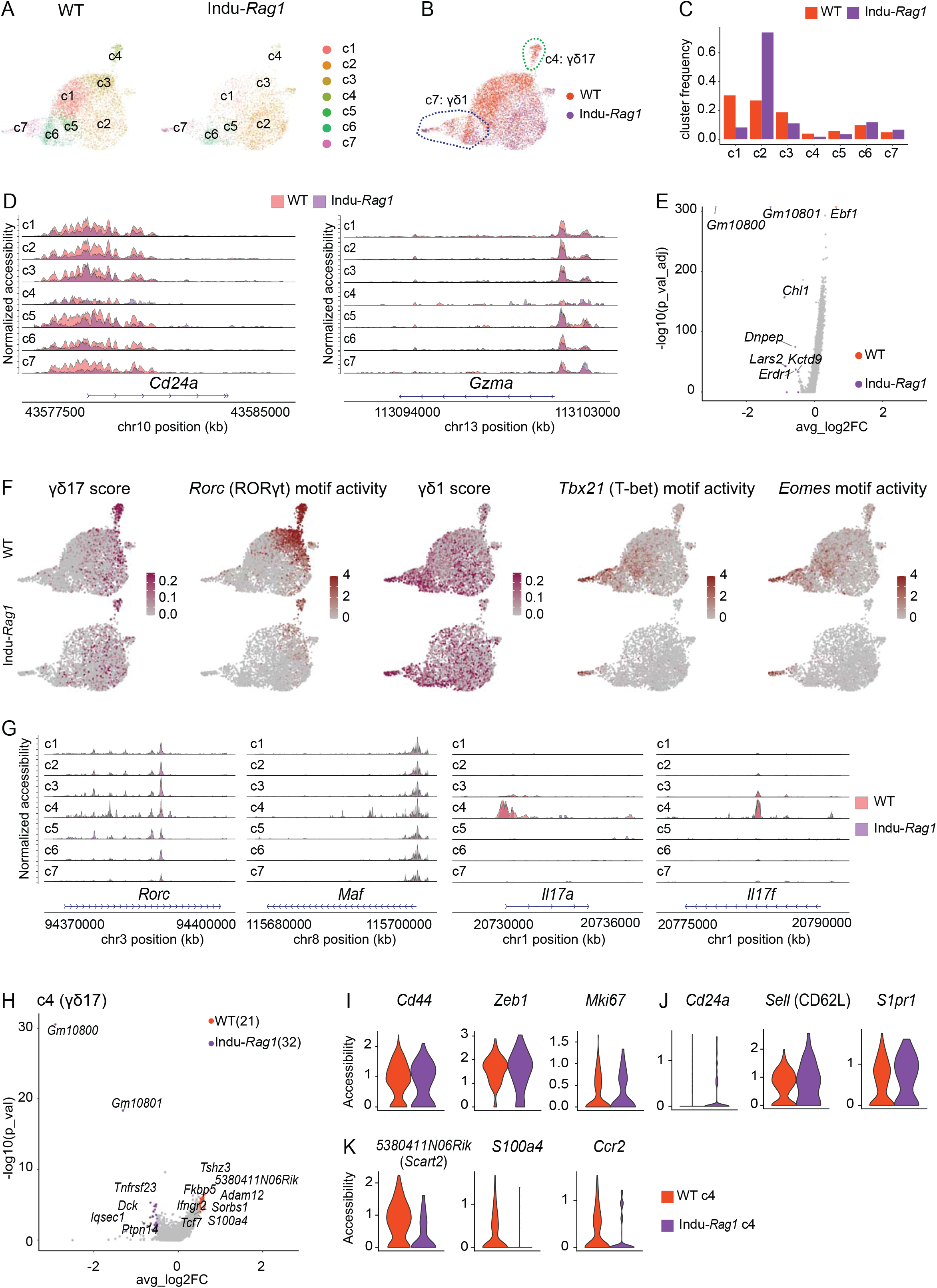
Vγ4 cell development from adult thymus precursors does not affect epigenetic programs in thymus. (**A-B**) Umap representation of WT and Indu-*Rag1* Vγ4 cell scATACseq data, colored by Cluster (**A**) and sample (**B**). Indu-*Rag1* thymus and LN Vγ4 cells were isolated by flow cytometric sorting from six adult mice. (**C**) The bar plot reveals fractions of absolute cell numbers from WT and Indu-*Rag1* Vγ4 cells that contributed to c1 to c7. (**D**) Genome track visualization of the *Cd24a* and *Gzma* locus of each Cluster in WT and Indu-*Rag1* Vγ4 cells, colored by sample. (**E**) Volcano plot demonstrates DAGs of WT and Indu-*Rag1* Vγ4 cells. Upregulated DAGs are identified with log2FC > 0.5 and p_val_adj < 0.05. (**F**) Umap shows the γ*δ*17 accessibility, *Rorc* motif activity, γ*δ*1 accessibility, *Tbx21* and *Eomes* motif activity score. Among that, γ*δ*17 and γ*δ*1 accessibility score were computed with the same gene sets as in Fig. 1E. (**G**) Genome track visualization of the *Rorc, Maf, Il17a* and *Il17f* locus of each Cluster in WT and Indu-*Rag1* Vγ4 cells, colored by sample. (**H**) Volcano plot demonstrates DAGs of WT and Indu-*Rag1* γ*δ*17 Vγ4 Cluster (c4). Upregulated DAGs are identified with log2FC > 0.5 and p_valj < 0.05. (**I-K**) Violin plots show selected marker genes accessibility of WT and Indu-*Rag1* γ*δ*17 Vγ4 Cluster (c4), colored by sample.

A reduction of IL17A- and IL17F-producing γ*δ* T cells was observed in Indu-*Rag1* thymus (**Fig. 5**). Thus, we next asked about potential differences in molecular programs of γ*δ*17 cell Cluster c4. Notably, WT and Indu-*Rag1* thymus Vγ4 cells had similar *Il17a, Il17f, Maf* or *Rorc* gene accessibility (**Fig. 6G**). Moreover, only few differentially accessible gene regions among γ*δ*17 cells (Cluster c4) of Indu-*Rag1* and WT thymus were identified (**Fig. 6H**). Similar accessibility of effector marker *Cd44*, TCR signaling repressor TF *Zeb1*, and proliferation marker *Mki67* were evident (**Fig. 6I**). The immature marker *Cd24a, Sell* (encoding CD62L) and *S1pr1* display a slightly higher accessibility in Indu-*Rag1* conditions (**Fig. 6J**). WT-derived γ*δ*17 cells in c4 show higher gene accessibility of *Scart2, S100a4*, and *Ccr2* than Indu-*Rag1* (**Fig. 6K**). Importantly, *Scart2* encoding a member of the Scart scavenger receptor family, which was described to be expressed on perinatal-derived IL-17^+^ cells,^35^ was completely closed in Indu-*Rag1* thymocytes.

Together, the Indu-*Rag1* mouse model provides evidence for parallel epigenetic programs of thymus Vγ4 cell subsets upon *de novo* generation from adult thymus precursor cells. It further identified potential key molecules such as Scart2 that may serve as a marker and/or drive the generation of CD44^high^CD45RB^neg^ Vγ4 cells during the development in the embryonic thymus.

## Discussion

In this study, single-cell NGS approaches deciphered genetic programs of Vγ4 cells in adult thymus and LN, and after induction of T cell development in adult thymus precursor cells. In contrast to previous multi-omics studies,^36^ the generated high resolution map of single-cell transcriptional and epigenetic profiles can be clearly assigned to Vγ4 cells. This provides a valid resource to define developmental and effector stages of Vγ4 T cells, including the identification of thus far undescribed regulators of γ*δ* T cell maturation. For example, the transcription factor Zeb1 was counter acting the expression of molecules related to TCR signaling of thymic γ*δ*24 cells (e.g. *Lck*). During the development of unconventional T cells, *Zeb1* represses TCR signaling and promotes cell proliferation. Such functional deficiency results in loss of peripheral Nk1.1^+^ γ*δ* T cells.^37^ The gene expression of *Zeb1* within thymic Vγ4 cells observed here supports the idea that strength of Vγ4^+^ TCR signaling is tightly regulated. ^38–40^ Future studies need to address the role of Zeb1 during γ*δ* T cell development and it’s restriction to Vγ4 cells.

It is perceived that IL-17-producing γ*δ* T cells that have highly invariant TCRs composed of either the Vγ6 or Vγ4 γ-chain do exclusively derive from the embryonic thymus.^6,11^ In contrast, few studies suggest that IL-17 production can be induced also from postnatal thymus-derived Vγ4 cells under certain circumstances.^20,21,32^ Thus, it remains controversial whether Vγ4^+^ γ*δ*17 cells derive from the pre-as well as the postnatal thymus. Therefore, we intended to re-investigate transcriptional programs of developing Vγ4 cells in adult thymus. We also employed a mouse model where the intentional activation of *Rag1* leads to the generation of exclusively postnatal thymic Vγ4 cells.

Our NGS analysis confirmed the presence of *Cd24a*^+^ and *Gzma*^+^ Vγ4 cells, which were recently identified within total γ*δ* T cells of fetal and adult thymus.^13,16,41^ As their granzyme A production capability never reached the level of NK1.1^+^ αβ T cells, we suggest a default pre-programming towards *Gzma*^+^ expression that is immediately lost upon further functional maturation. The *Gzma*^+^ γ*δ* T cells do also express c-Maf. c-Maf is a universal TF of IL-17-producers.^14^ This highlights that developing *Gzma*^+^ cells harbor molecular programs of γ*δ*17 T cells in adult thymus. Consistent with this idea a small, but considerable fraction of developing CD24^+^ Vγ4 cells that has IL-17 production capability in adult thymus was observed.

Employing a model where tamoxifen-induced *Rag1* activation leads to T cell development in adult thymus, we revealed that developmental plasticity is independent of the ontogenetic timing. Evidence is also provided for the *de novo* generation of adult thymus derived c-Maf^+^ and RORγt^+^ Vγ4 cells. However, these *de novo* generated γ*δ*17 T cells did not reach the periphery. Mature CD44^high^CD45RB^neg^ Vγ4 cells were completely absent within all studied Indu-*Rag1* samples. This implies that specific signals for the full maturation and thymic export of cMaf^+^ RORγt^+^ Vγ4 cells are missing in adult thymus. Instead, in normal mice the large majority of LN (and thymus) Vγ4 cells express the naïve T cell marker CD62L. This surface molecule is implied in the activation and recruitment of γ*δ* T cells into inflamed tissues.^42^ Its expression was found to be similar on Vγ1 cells. In line with the absence of innate effectors in Indu-*Rag1* samples, we found a slightly higher gene accessibility of *Sell* (encoding CD62L) in the Indu-*Rag1 Cd24a*^+^ γ*δ*17-like ATACseq Cluster as compared to WT conditions. It was previously reported that antigen-stimulation can induce the differentiation of CD62L^+^ γ*δ* T cells into IL-17-producing effector cells.^43^ future studies need to investigate whether adult-thymus derived Vγ4 cells retained the potential to become functional IL-17 producers during infection or in the onset of inflammatory diseases.

Short- and long-time *ex vivo* stimulation with the innate-cytokines IL-1ß and IL-23 confirmed the impaired capability of adult-derived γ*δ*17-like cells to produce IL17-A and IL17-F production. In contrast to an alternative genetic mouse model where cells derive from different precursor cell stages, IL-17A-producing γ*δ* T cells were hardly detectable in the lung^32^ and lymph nodes of Indu-*Rag1* mice. Therefore, we propose that adult thymus derived γ*δ*17-like cells lose their phenotype. They might undergo apoptosis or simply follow developmental traits that bias them towards a higher generation of naïve-like cells than fetal immature γ*δ*17 cells. The development and/or final differentiation of these γ*δ*17-like cells may also depend on differential requirements for Notch signaling. Notch is known to be one of the key factors for γ*δ*17 cell fate decisions in the perinatal period.^19,32^ Alternatively, they might be derived from different precursors or/and may proceed via different developmental pathways compared to perinatal cells.^18^

To our surprise, only very few differentially accessible gene regions were identified within WT and Indu-*Rag1* γ*δ*17 cell Clusters. On the other hand, NGS profiling revealed that the gene locus for Scart2, a protein associated with IL-17-producing Vγ4 T cells,^35,44^ was completely closed within the adult Indu-*Rag1* thymus. These data imply that Scart2 is differentially expressed only on Vγ4^+^ γ*δ*17 cells of embryonic and perinatal origin which are believed to be functionally pre-committed towards IL-17 production already before TCR rearrangement.^6,35^ Interestingly, Scart1 another family member can be employed as a marker for fetus-derived Vγ6 cells.^9,44^ This poses the question on the involvement of Scart1 or Scart2 proteins on IL-17 effector fate decisions. Future studies need to invest in their involvement in fine-tuning TCR^44,45^ and/or Notch^19^ signals when the cells undergo final differentiation during thymic development, including the identification of fetal versus adult thymus-derived signals to guide effector pre-programming.

Recent NGS studies reported the generation of human γ*δ* T cells with transcriptional profiles of γ*δ*17 cells within the early human embryonic thymus.^46,47^ This highlights similarities in the ontogeny of γ*δ*17 cells between mice and men. At the same time, a small proportion of γ*δ*17-like T cells was identified within the postnatal human thymus. However, unlike the human fetal γ*δ*17-like T cells, these cells from pediatric thymi did not express *Il17a* transcripts.^46^ Again raising the question on the function and relevance of IL-17 producing γ*δ* T cells derived from the early *versus* adult life period. The adult-thymus derived c-Maf^+^ RORγt^+^ Vγ4 cells described here do not complete full maturation into peripheral IL-17 producers.They may represent murine counterparts of the γ*δ*17-like cells identified in the pediatric human thymus. As the newly generated murine Vγ4 cells reached some γ*δ*17 functions within the adult thymus. Future studies need to solve the question about potential signals that inhibit the full development of adult thymus-derived γ*δ*17 cells and how they perform under inflammatory conditions.

## Methods

### Mice

The C57BL/6NCrl, C57BL/6-Trdc^tm1Mal^ (here Tcrd-H2BeGFP)^48^ and B6-Trcdtm1Mal Rag1tm1.1Sadu Gt(ROSA)26Sortm1(creERT2)Tyj (here *Indu-Rag1*)^6^ mice were used in the study. Animals were housed under specific pathogen free conditions at the central animal facility of Hannover Medical School, Germany.

For all experiments, adult mice of both gender were used and allocated randomly to the experiment. For dissection of organs and preparation of single cell suspensions mice were sacrificed by CO2-inhalation and cervical dislocation. For 5-6 weeks old Indu-*Rag1* mice, conditional *Rag1* expression was induced at 5-6 weeks old by oral supply of 400 µl (4 mg/400 µl) tamoxifen citrate (Enzo Life Sciences, dissolved in corn oil/ethanol). Three or six weeks after *Rag1* induction cells were harvested for flow cytometric analysis and scATACseq. All experimental procedures were conducted according to institutional guidelines approved by Lower Saxony State Office for Consumer Protection and Food safety animal care and use committee (reference number 17/2704 and 2021/276).

### Isolation of lymphocytes

Single cell suspensions of thymus and pLNs cells (including superficial cervical, axillary, brachial and inguinal LNs) were mashed and filtered with nylon gaze. To enrich γ*δ* T cells in thymus samples, CD4+ and CD8+ T cells were lysed as described before.^6^ Briefly, cells were incubated with monoclonal anti-CD4 and anti-CD8b IgM (homemade), DNaseI (Roche) and Standard Rabbit Complement (Cederlane) at 37°C for 30 minutes. Next, cells were retrieved via Lympholyte M (Cederlane) density gradient centrifugation and re-suspended in MACS buffer (PBS with 3% FCS and 4mM EDTA).

The isolation of lymphocytes from ear skin was done as described before.^9^ Briefly, ears were split, cut, and incubated with digestion buffer (RPMI media supplemented with (2mg/ml) collagenase IV (Worthington) and (187.5 μg/ml) DNaseI (Roche)) for 75 minutes at 37°C under 1400 rpm of shaking. After stopping the digestion with 5mM EDTA for 15 minutes at 37°C under 1400 rpm of shaking, ear tissue was further dissociated via a 18G needle. Next, cells were filtered through a 100 μM cell strainer (Falcon) for density gradient centrifugation using 40% and 70% Percoll solutions and re-suspended in MACS buffer.

To isolate lymphocytes from lung, lungs were cut into small pieces, and digested with digestion buffer (0.5 mg/ml Collagenase D (Worthington) and 0.025 mg/ml DNase I) for 45 minutes at 37°C. After stopping the digestion with 20mM EDTA, lung tissue was then filtered with 40 µm pore Cellstrainer. Cells were retrieved via Lympholyte M density gradient centrifugation and re-suspended in MACS buffer.

### Flow cytometry

After obtaining the single cells suspension, cells were blocked with FcR antibody (clone 2.4G2) on ice for 5 min, followed with a cocktail of surface antibodies at optimized concentrations for 20 min on ice. The Vγ6 staining was done as described.^49^ Briefly, the cells were stained with IgM anti-Vγ5/6 after preincubation with unlabeled anti-Tcrd (clone GL3, homemade) and other surface antibodies. Afterward, the Vγ6 cells were detected with anti-IgM PE (clone RM-7B4, eBioscience).

For GZMA/RORγt/c-Maf intracellular staining, cells were fixed and permeabilized with Foxp3 staining kits (eBioscience), according to the manufacturer’s protocol. For IL-17A/IL-17F/IFN-γ intracellular staining, the ICS staining buffer set (eBioscience) was used. Following antibodies were employed (clone, company): anti-TCRβ APCVio770 (REA318, Miltenyi), anti-TCRβ PE-Cy5 (H57-597, BD), anti-CD3 APC-Fire810 (17A2, BioLegend), anti-Vγ4 BV480 (UC3-10A6, BD), anti-Vγ4 APC (UC3-10A6, BioLegend), anti-Vγ1.1 BV711 (2.1, BD), anti-Vγ1.1 APC (2.1, BioLegend), anti-CD44 BUV496 (IM7, BD), anti-CD45RB (126A, BD) anti-CD24 PE (M1/69, BioLegend), anti-CD27 BUV563 (LG.3A10, BD) anti-CD73 BV605 (TY/11.8, BD), anti-CD62L BV711 (MEL-14, BioLegend), anti-c-Maf (eFluor660, Thermo Fisher), anti-RORγt PerCP-ef710 (B2D, Thermo Fisher), anti-GzmA eFluor450 (GzA-3G8.5, Thermo Fisher), anti-IL-17F PerCP-ef710 (SHLR17, Thermo Fisher), anti-IFN-γ PE-Cy7 (XMG1.2, BD) and anti-IL-17A APC (TC11-18H10.1, BioLegend). Zombie Aqua and Zombie NIR Fixable Viability Kit (BioLegend) were employed for dead cell exclusion. Flow cytometry was performed using BD LSRII (BD) equipped with three lasers operating on 405nm, 488nm and 640nm using BD FACS diva () or Cytek Aurora spectral flow cytometer (Cytek) equipped with five lasers operating on 355nm, 405nm, 488nm, 561nm and 640nm using SpectroFlo v2.2.0 (Cytek). Data analysis was performed using Flowjo 10.0 software or FCS Express V7 (Denovo).

Sorting was performed in the Cell Sorting Core Facility of the Hannover Medical School by using the FACSAria Fusion (BD). For all NGS experiments Vγ4 cells were FACS sorted as Zombie Aqua^-^GFP^+^ TCRβ^-^ Vγ4^+^ lymphocytes.

### Ex vivo cell stimulation

For ex vivo stimulation, single cell suspensions were either enriched for γ*δ* T cells using Mojo Sort magnetic beads (BioLegend) for depletion of CD4 and CD8 T cells or sorter for Vγ4 T cells. Cells were incubated with complete medium in the presence of murine IL-1β (10 ng/ml) and IL-23 (10 ng/ml) for the indicated duration. Next, cells were re-stimulated with PMA (50 ng/ml) and ionomycin (2 µg/ml) and incubated for 4 h at 37 °C with Brefeldin A (10 µg/ml), followed by flow cytometric analysis.

### scRNAseq library and bioinformatics flow

Isolated thymus and pLNs Vγ4 cells were directly subjected to scRNAseq library construction using the Chromium Next GEM Single Cell V(D)J Reagent Kits v1.1 according to the manufactures protocol (10xGenomics). Libraries were sequenced with Illumina NexSeq 500 platform. Sequence reads were aligned to reference mouse genome mm10 (UCSC) and cell barcode-gene matrices were counted with Cell Ranger 3.1 (10x Genomics). The cell barcode-gene matrices were then processed with Seurat v4.0.1 under R v4.0.3 to remove low quality cells (genes > 200, Features < 4000, %Mitochondrial genes <20).^50^ Thymus and LN Vγ4 cell scRNAseq data were merged and performed the further normalization, scaling, and dimension reduction using functions from Seurat. The batch effect was removed with function “RunHarmony”, a Seurat function applied from Harmony package.^51^ Cells were clustered with “FindCluster” function and annotated according to the expression of lineage-specific markers (visualized with “DotPlot” function). Aggregated lineage marker expression scores were calculated with “AddModuleScore” function, and then visualized with “FeaturePlot” function. γ*δ*1 and γ*δ*17 subsets were isolated with “subset” function by choosing the respect Clusters. Differential expression gene (DEG) analysis between organs was assessed with “FindMarkers” function for transcripts detected in at least 10% of cells using logFC (log-fold-change) threshold of 0.5.

### scATACseq library and bioinformatics flow

Nuclei of thymus and pLN Vγ4 cells isolated from WT or Indu-*Rag1* animals were isolated according to the protocol provided by 10xGenomics. Briefly, the cells were washed with 0.04% BSA PBS buffer and then incubated in a lysis buffer (10 mM Tris-HCl, 10 mM NaCl, 3 mM MgCl2, 0.1% Tween-20, 0.1% NP40, 0.01% digitonin and 1% BSA in nuclease-free water). After 3 min lysis, the nuclei were washed with buffer (10 mM Tris-HCl, 10 mM NaCl, 3 mM MgCl2, and 0.1% Tween-20) and re-suspended in Nuclei Buffer (10xGenomics), followed by immediate processing for transposition and library construction using Chromium Next GEM Single Cell ATAC Reagent Kits v1.1 according manufactures protocol (10xGenomics).

Libraries were sequenced at the Illumina NexSeq 500 platform. Sequence reads were aligned to reference mouse genome mm10 (UCSC) and cell barcode-peaks matrices were counted with Cell Ranger ATAC 1.2 (10x Genomics). Subsequently analysis was performed with Signac 1.3.0, a package for scATACseq analysis derived from Seurat.^52^ Low-quality cells were removed (peak region fragments > 300, peak region fragments < 60000, % reads in peaks > 65 nucleosome signal < 1.2 & TSS enrichment score > 2.5) before normalization with term-frequency inverse-document-frequency (TFIDF) and dimension reduction for visualization using UMAP. Integrated analysis was performed with “merge” function followed by removing batch effect with “RunHarmony” function. Gene activity/accessibility matrix was calculated by counting peaks within the gene coding region and promoter region (2kb upstream region of the gene transcriptional start site). Cells were Clustered with “FindCluster” function and annotated according to the accessibility of lineage-specific markers (visualized with “DotPlot” function). Aggregated lineage marker accessibility scores were calculated with “AddModuleScore” function, and then visualized with “FeaturePlot” function. Differential accessible region/gene (DAR/DAG) analysis was performed with “FindMarkers” function for fragment detected in at least 10% of cells using logFC (log-fold-change) threshold of 0.5.

### Transcription factor motif analysis

Using functions “AddMotifs” and “RunChromVAR” from Signac and chromVAR,^53^ transcription factor motif activity matrix (motif from JASPAR version 2020) was obtained by counting the probability of the specific motif at the given frequency by chance comparing with a background set of peaks matched for GC content. TF binding to DNA protects the binding of Tn5 transposase creates increased DNA accessibility in the immediate flanking sequence. Hence, transcription factor motif footprinting was estimated using “Footprint” function.

### Pseudotime trajectory analysis

We assessed the pseudo-time trajectories with Monocle 3 and Cicero.^54,55^ The “as.cell_data_set” function was used to convert Seurat object into a cell dataset object (cds). The dataset was ordered by choosing c2, the highest *Cd24a* accessibility, as “start point” using “order_cell” function. The data of gene accessibility across pseudo-time were visualized with “plot_genes_in_pseudo-time” function.

### NGS data accessibility

Raw single-cell sequencing data is available under NCBIs Gene Expression Omnibus (GSE222588).

## Acknowledgments

We thank Matthias Ballmaier of the central cell sorting facility and the Genomics platform of the Hannover Medical School for support. We highly thank Dr. Likai Tan for scientific and technical advice. The study was supported by the German Research Foundation - Deutsche Forschungsgesellschaft (DFG) under Germany’s Excellence Strategy, EXC 2155 RESIST, Project ID 390874280 to I.P., R.F. and S.R.; the SFB900 Project ID 158989968 to I.P., R.F., and S.R.; and the DFG-funded research group FOR2799 Project ID RA3077/1-2 to S.R.

## Authors contributions

T.Y., Z.W., A. J. X.L. and J.BM. performed experiments; S.W., I.P. and R.F. contributed with resources; T.Y. analyzed data; S.R. wrote the manuscript; T.Y., J.BM. and S.R. designed research, experiments and wrote the first manuscript draft.

