## Supplement_Figures for "Adult thymus-derived cMaf^+^RORγt^+^ γδ T cells lack Scart2 chromatin accessibility and do not reach periphery"

### **Supplemental Figure. 1** Related to Figure. 1

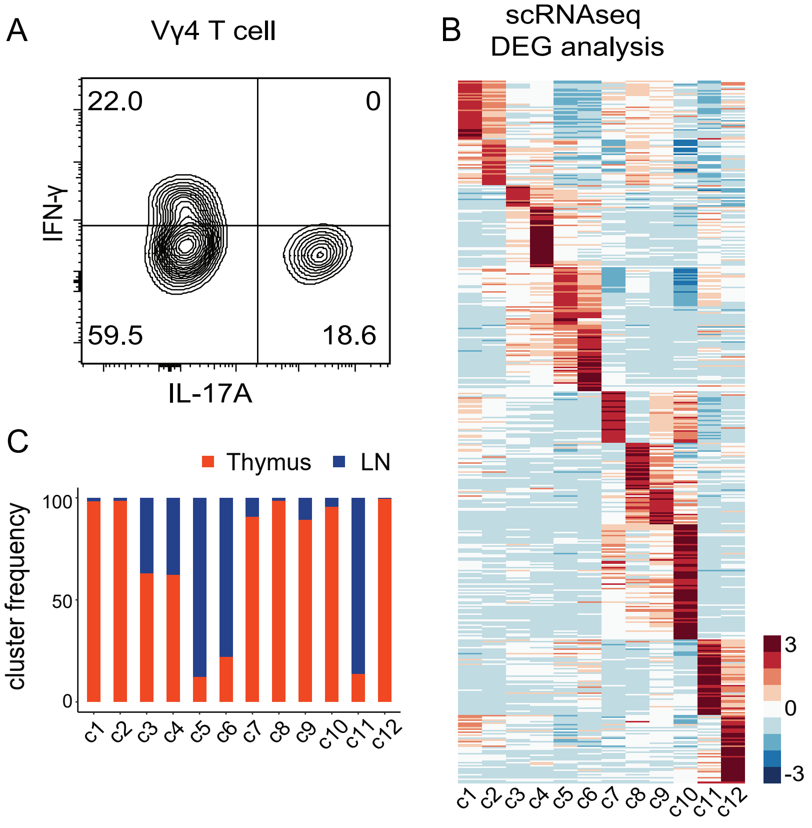


**Fig. S1: Quality control of scRNAseq data.** (**A**) FACS analysis of IL-17A and IFN-γ production in LN Vγ4 cells after stimulation with PMA plus ionomycin for 4h with Brefeldin A. (**B**) Heatmap represents the top 50 DEGs for each cell Cluster. (**C**) The bar plot reveals fractions of absolute cell numbers from thymus and LN Vγ4 scRNAseq data that contribute to c1 to c12.

**Supplemental Figure. 2** Related to Figure. 2

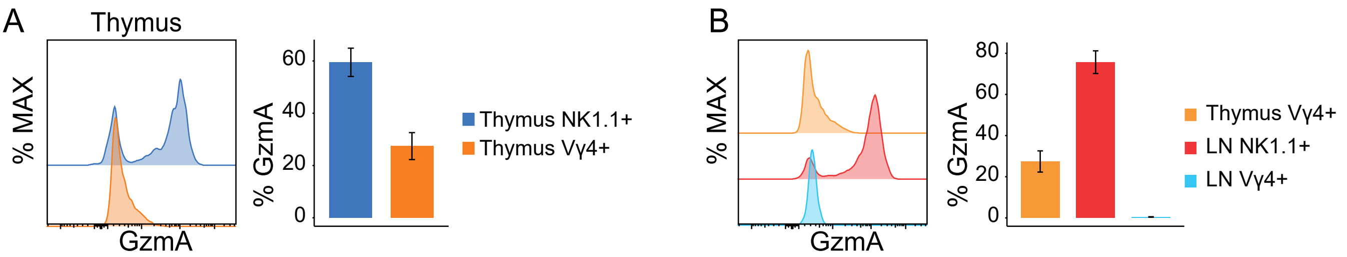


**Fig. S2: Characterization of GzmA$^{+}$ CD24$^{+}$ Vγ4 cells by flow cytometry.** (**A-B**) FACS plots and graph show the GZMA expression in thymus NK1.1$^{+}$ αβ T cells and Vγ4$^{+}$ T cells (N=3) (**A**), and in thymus Vγ4$^{+}$ T cells and LN NK1.1$^{+}$ and Vγ4$^{+}$ cells (N=3) (**B**).

**Supplemental Figure. 3** Related to Figure. 3

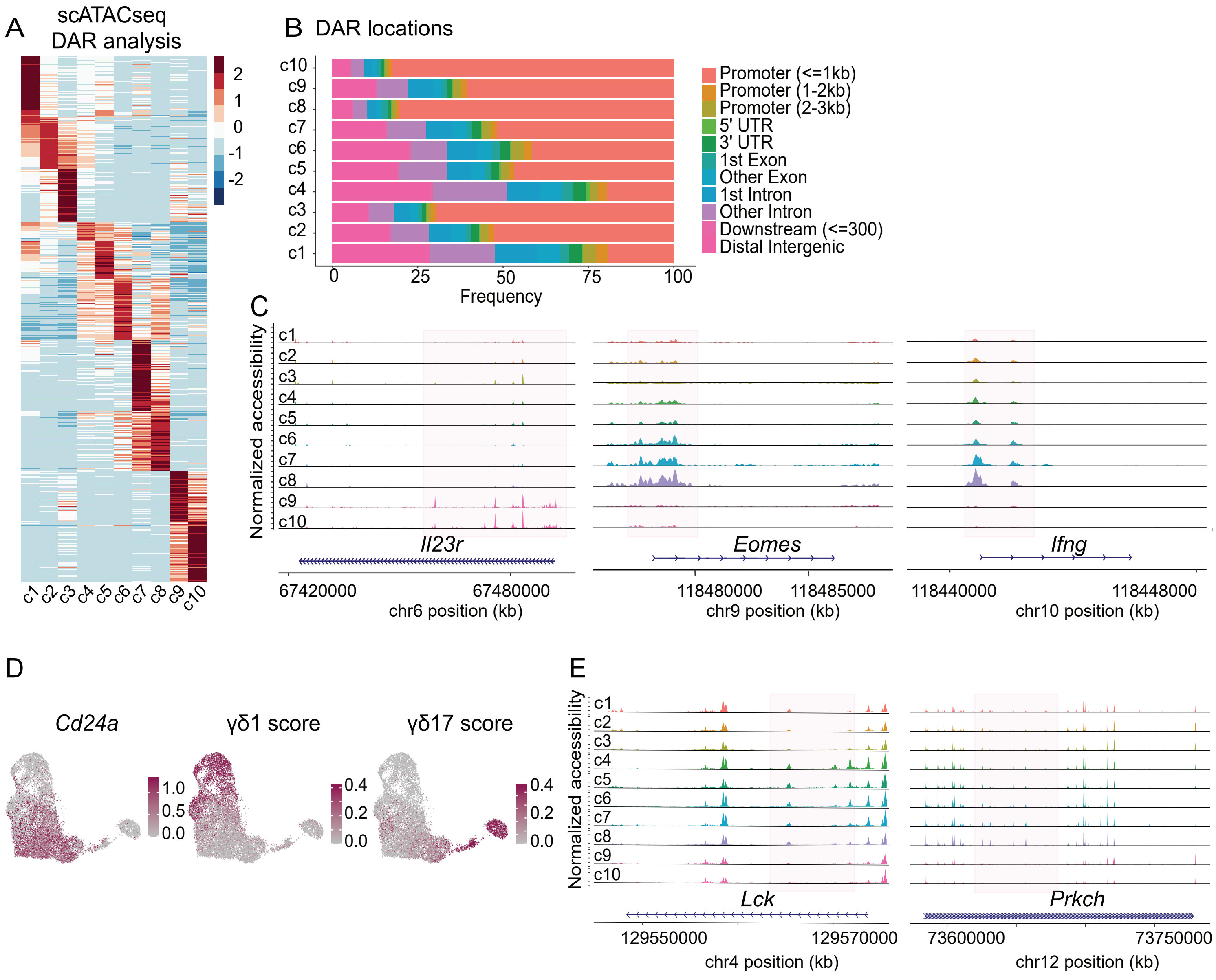


**Fig. S3: Quality control of scATACseq data.** (**A**) Heatmap represents the top 50 DARs for each cell Cluster. (**B**) Bar plot of annotated differential accessible regions (DARs) location of each Cluster. (**C, E**) Genome track visualization of the *Il23r*, *Eomes* and *Ifng* (**C**), and *Lck* and *Prkch* (**E**) locus of each Cluster. (**D**) Umap displays the *Cd24a* accessibility, γδ1 accessibility and γδ17 accessibility scores of each cell, computed with the same sets of genes with scRNAseq.

**Supplemental Figure. 4** Related to Figure. 3

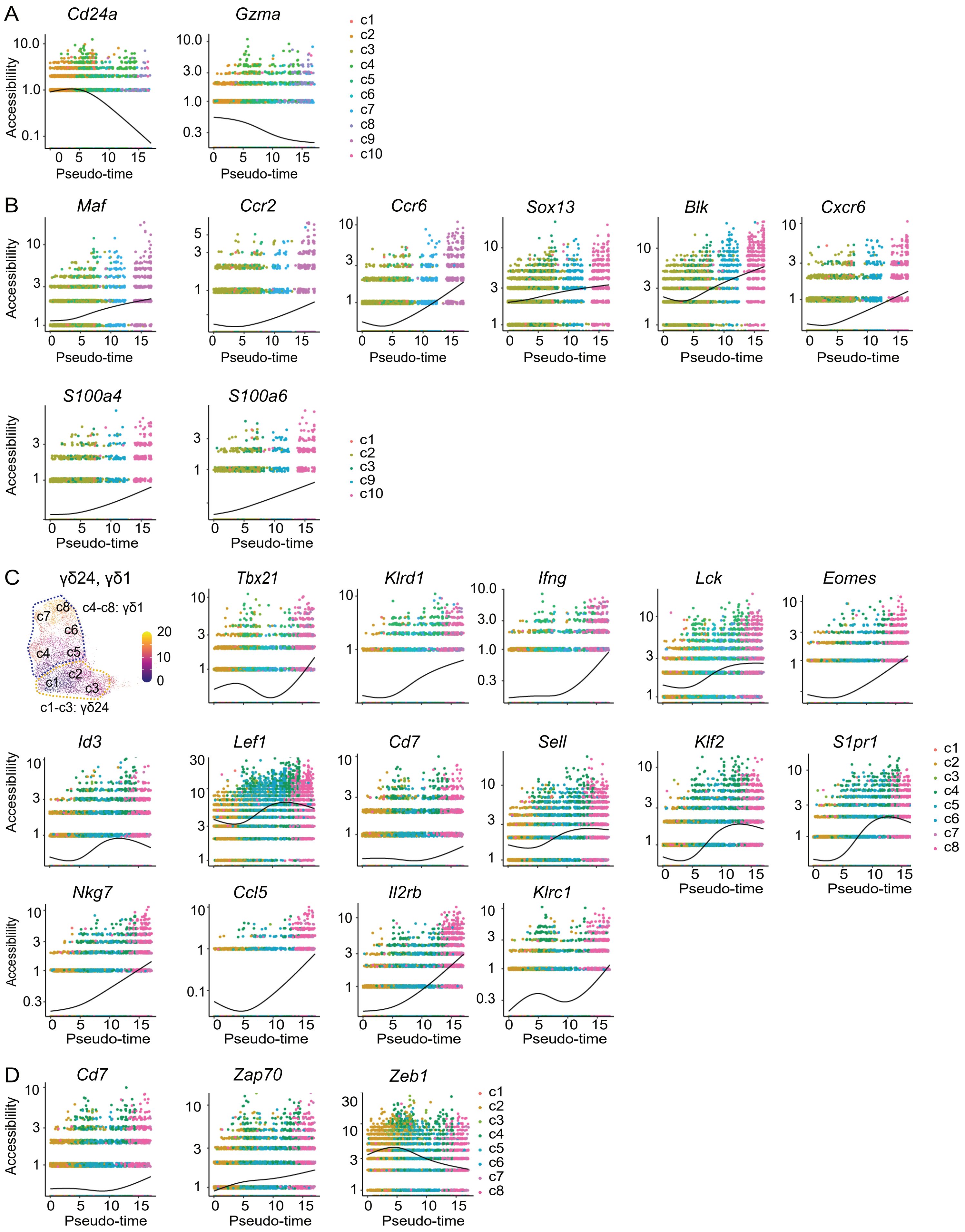

**Fig. S4: Pseudo-time analysis of Vγ4 cells in adult thymus.** (**A**) *Cd24a* and *Gzma* accessibility dynamics across pseudo-time in the total dataset. (**B**) Gene accessibility across the γδ17 pseudo-time trajectory: γδ24 (c1-c3) to γδ17 (c9-c10). (**C**) Umap displaying the pseudo-time trajectory from the γδ1 trait: γδ24 (c1-c3) to γδ1 (c4-c8) and gene accessibility dynamics across the pseudo-time trajectory. (**D**) Gene accessibility of TCR signaling marker genes (*Cd7*, *Zap70*, and repressor *Zeb1*) across the γδ1 trait: γδ24 (c1-c3) to γδ1 (c4-c8).

**Supplemental Figure. 5** Rerelated to Figure. 4

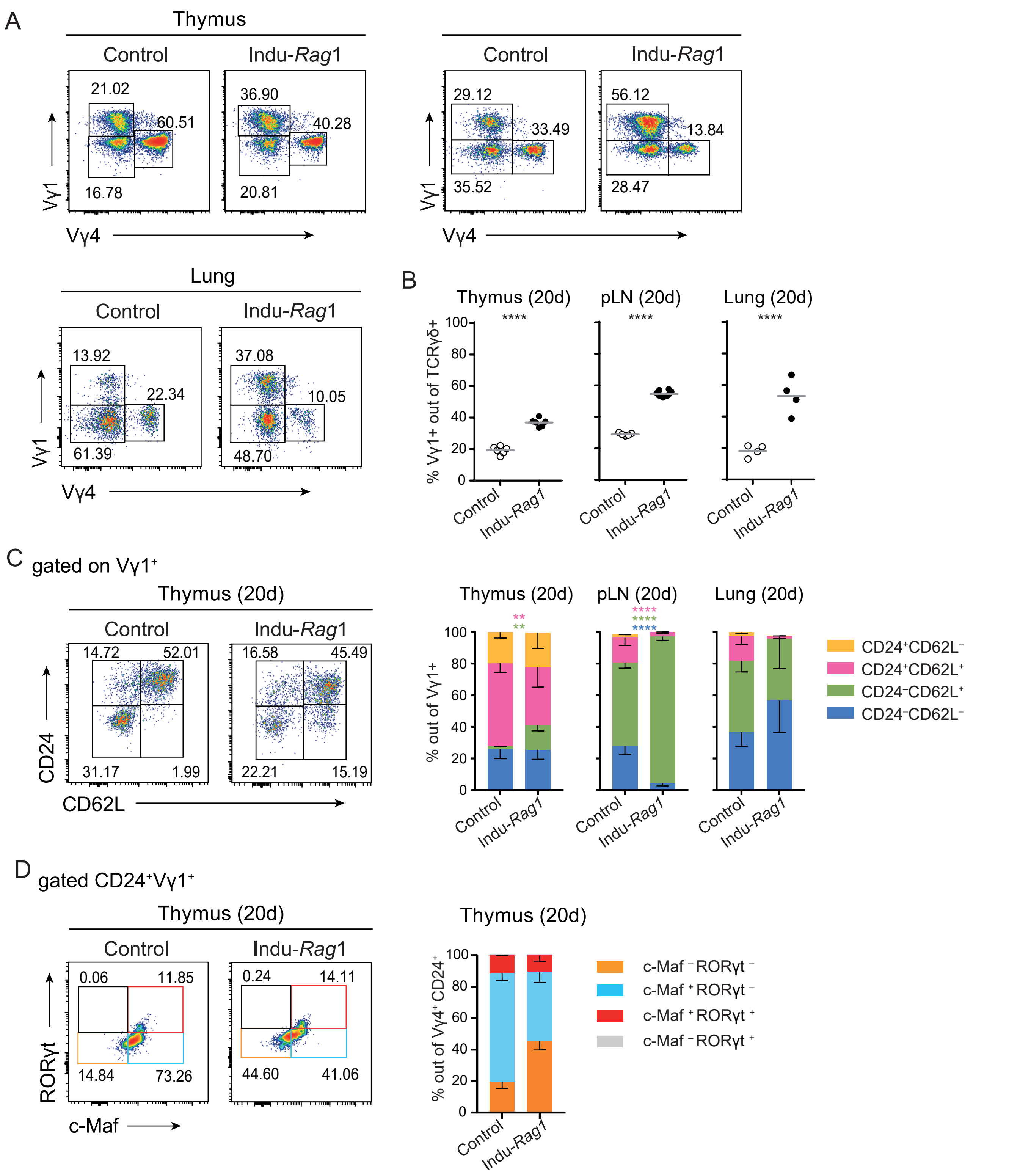


**Fig. S5: Indu-*Rag1* mice lack Vγ6⁺ γδ T cells but can generate Vγ1⁺ with identical traits to controls.** Gating strategy (**A**) and quantification (**B**) of flow cytometry analysis of Vγ chain usage in thymus, peripheral Lymph Nodes (pLN) and Lung of 20 days old controls and Indu-*Rag1* mice 20 days post induction of the *Rag1* gene. (**C**) Gating strategy and quantification of flow cytometry analysis of CD24 and CD62L expression in Vγ1 γδ T cells from thymus (left graph), pLNs (center graph) and lung (right graph) of 20 days old control mice and Indu-*Rag1* 20 days after induction. (D) Gating strategy (left) and frequency distribution (right) of RORγt and c-Maf expressing populations out of CD24⁺Vγ1⁺ in 20 days old control mice and Indu-*Rag1* mice 20 days post induction.

### **Supplemental Figure. 6** Rerelated to Figure. 6

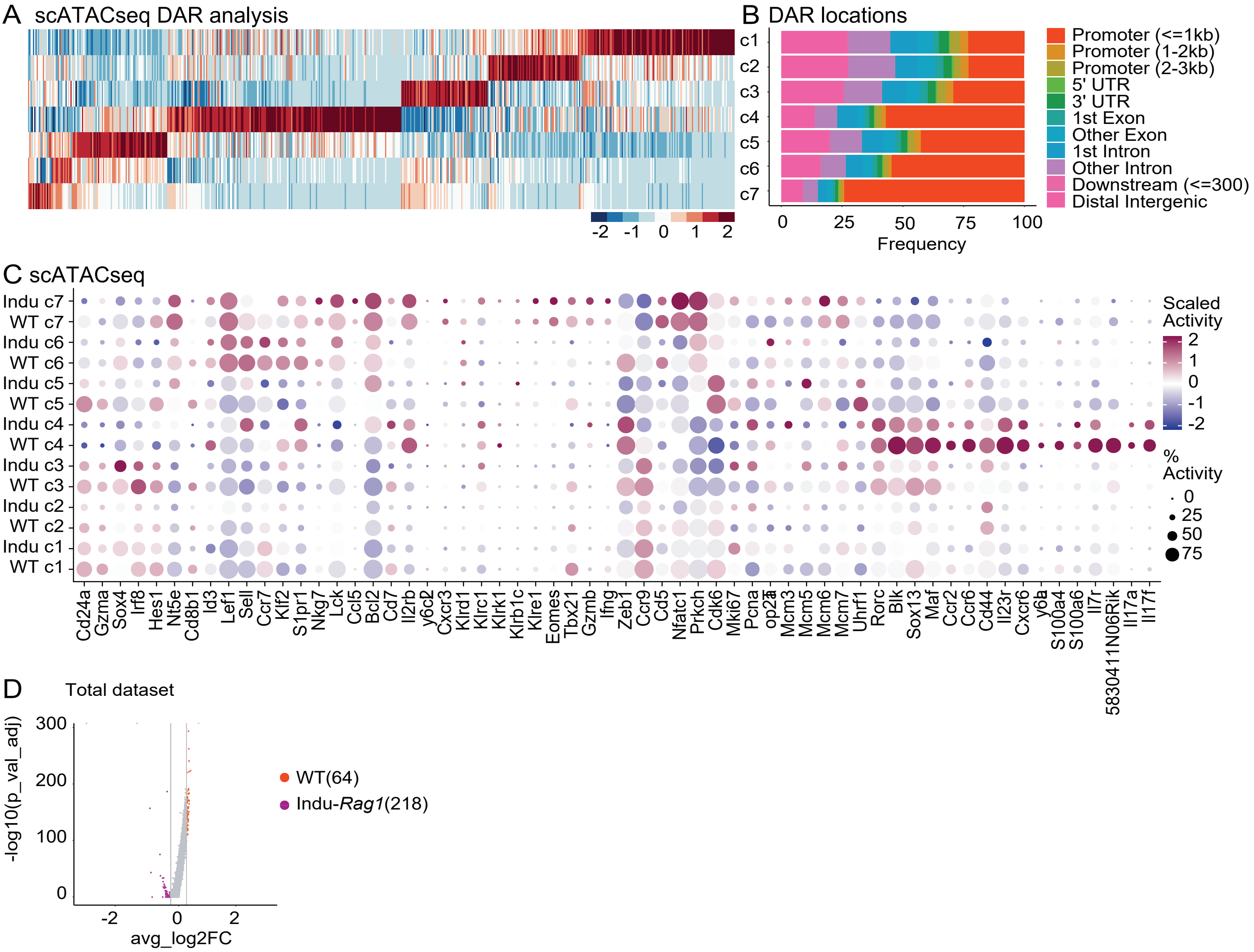


**Fig. S6: Comparison of epigenetic profiles of Vγ4 cells in WT and Indu-*Rag1* thymus.** (**A**) Heatmap represents the top 50 DARs for each cell Cluster. (**B**) Bar plot of annotated DARs location of each Cluster. (**C**) Dot plots showing the selected up-regulated marker DAGs (gene body ± 2 kb) of each Cluster in each sample. (**D**) Volcano plot demonstrates DAGs of WT and Indu-*Rag1* Vγ4 cells. Upregulated DAGs are identified with log2FC > 0.25 and p_val_adj < 0.05.
